# Soluble CD155 immune complexes reprogram DNAM-1 signaling to potentiate antitumor immunity

**DOI:** 10.64898/2026.05.16.725599

**Authors:** Shota Kinoshita, Tomohei Matsuo, Naoto Takeuchi, Genki Okumura, Natsuki Ide, Akiko Iguchi-Manaka, Soontae Gwon, Eri Sugiyama, Shohei Koyama, Hideaki Tahara, Chigusa Nakahashi-Oda, Akira Shibuya, Kazuko Shibuya

## Abstract

Resistance to immune checkpoint inhibitors (ICIs) remains a major challenge in oncology, yet the mechanisms that selectively disable activating pathways are poorly defined. Here, we identify tumor-derived soluble CD155 (sCD155) as a systemic checkpoint that rewires DNAM-1 signaling to drive immunotherapy resistance. High plasma sCD155 levels correlate with impaired anti-PD-1 responses in patients with non-small cell lung cancer. Mechanistically, sCD155 selectively suppresses DNAM-1-dependent activation of CD8^+^ T and NK cells, uncoupling ICIs from cytotoxic function. Intriguingly, a selective anti-sCD155 monoclonal antibody does not neutralize the ligand, but rather converts it into an activating scaffold. This complex induces FcγR-anchored DNAM-1 microcluster formation and robust downstream signaling, effectively switching the checkpoint into a co-stimulatory signal. This reprogramming restores CTL function, suppresses metastasis, and augments PD-1/TIGIT blockade to achieve durable immunity. Our findings establish antibody-mediated receptor architecture rewiring as a therapeutic principle to overcome cancer immune resistance.

## Introduction

Immune checkpoint inhibitors (ICIs), most prominently antibodies targeting programmed cell death protein 1 (PD-1) and its ligand PD-L1, have reshaped cancer therapy by unleashing antitumor T cell immunity and producing unprecedented clinical benefits across multiple malignancies (1,2). Despite their transformative impact, durable responses are achieved in only a minority of patients, underscoring the existence of additional immunoregulatory circuits that constrain therapeutic efficacy (3). Consequently, intensive efforts have focused on identifying cooperative checkpoint pathways that determine responsiveness to ICIs.

Among emerging regulators of antitumor immunity, the poliovirus receptor CD155 has attracted considerable attention as a central ligand that integrates activating and inhibitory immune signals within the tumor microenvironment (4–7). CD155 is a stress-inducible molecule frequently upregulated in diverse malignancies (8). Through engagement of multiple receptors—including the costimulatory receptor DNAM-1 (CD226) and the coinhibitory receptors T cell immunoreceptor with Ig and ITIM domains (TIGIT), CD96, and KIR2DL5—CD155 orchestrates a finely balanced signaling network that controls cytotoxic lymphocyte activity (9–12). Whereas TIGIT, CD96, and KIR2DL5 suppress effector functions of natural killer (NK) cells and CD8⁺ T cells, DNAM-1 provides a dominant activating signal essential for effective antitumor immunity and clinical responses to PD-1 blockade (13–18).

Recent studies have further revealed that DNAM-1 represents a pivotal node within immune checkpoint therapies. TIGIT not only competes for CD155 binding but actively suppresses DNAM-1 expression and signaling through post-translational mechanisms, including receptor internalization and degradation (19, 20). Consistently, DNAM-1 expression and phosphorylation status—rather than TIGIT levels—correlate with responsiveness to PD-1 blockade in patients with non-small cell lung cancer (7). These findings position DNAM-1 as a critical determinant of checkpoint therapy efficacy and suggest that perturbations of the CD155–DNAM-1 axis may fundamentally limit immunotherapeutic outcomes.

CD155 is strongly associated with tumor progression and poor clinical outcomes (21, 22). While several mechanisms have been proposed to explain this correlation, the precise processes remain to be fully elucidated. In humans, CD155 exists not only as membrane-bound isoforms (CD155α) but also as soluble splice variants (CD155β and CD155γ) lacking the transmembrane domain, collectively referred to as soluble CD155 (sCD155 (23)). Functionally, sCD155 suppresses NK cell-mediated cytotoxicity by interfering with DNAM-1 activation (24). Although sCD155 is known to interfere with the DNAM-1/TIGIT axis (24–27), its definitive role as a driver of poor prognosis through immune evasion has not been fully established. Moreover, its contribution to resistance against ICIs ̶and the precise mechanisms by which it rewires antitumor immune signaling—remains largely unexplored.

Here, we identify sCD155 as a previously unrecognized systemic immune checkpoint that actively limits the efficacy of cancer immunotherapy. Using tumor models and clinical analyses of patients with non-small cell lung cancer (NSCLC), we demonstrate that elevated sCD155 attenuates antitumor immunity and serves as a predictor of poor clinical responses to PD-1 blockade. To therapeutically target this axis, we developed a selective anti-sCD155 monoclonal antibody (mAb) that reprograms sCD155 into a DNAM-1 agonist via Fcγ receptor–mediated clustering. This strategy restores cytotoxic lymphocyte function and markedly enhances the efficacy of PD-1 and TIGIT blockade in vivo. Together, our findings reveal a soluble immune checkpoint–driven resistance mechanism and establish antibody-mediated rewiring of DNAM-1 signaling as a new principle for overcoming immunotherapy failure.

## Results

### NSCLC patients with higher sCD155 concentration showed reduced progression-free survival

To analyze the association between CD155 and sCD155 expression, we revisited the dataset on mRNA expression of PVRα, PVRβ, and PVRγ, which encode CD155 and sCD155, respectively, in 16 patients with colorectal, gastric, and breast cancers (26). We found that the relative expression of *PVR*α is strongly positively associated with that of *PVR*β and *PVR*γ (Figure S1A), indicating that tumors with high CD155 expression also release substantial amounts of sCD155 into the circulation. These results suggest that sCD155 might contribute to the poor prognosis of patients with tumors that express high levels of CD155.

To assess the clinical relevance of sCD155 for immune checkpoint inhibitor efficacy, we conducted a retrospective analysis of pretreatment plasma samples from patients with NSCLC. Plasma samples were collected from 49 patients prior to pembrolizumab therapy, administered either as monotherapy or in combination with chemotherapy (carboplatin plus pemetrexed or paclitaxel). Only patients who received at least two treatment cycles were included in the analysis. Baseline plasma sCD155 concentrations were quantified by ELISA, and patients were stratified into high and low groups using a cutoff value of 260 ng/mL (Figure 1A). Fourteen patients (28.6%) were classified as the high sCD155 group (mean 306.4 ng/mL, range 267.6–412.8), while 35 patients (71.4%) comprised the low sCD155 group (mean 189.2 ng/mL, range 112.8–256.7) (Figure 1A). No significant associations were observed between sCD155 levels and clinicopathological characteristics (Table S1). Kaplan–Meier analysis revealed that patients with high sCD155 levels had significantly shorter progression-free survival (PFS) than those with low sCD155 levels (Figure 1B). Consistent with previous reports (28), higher PDL-1 tumor proportion score (TPS) showed a trend toward improved PFS following pembrolizumab treatment (Figure S1B). Notably, however, within subgroups with tumors in which PDL-1 TPS is 1% or higher, patients with high sCD155 levels demonstrated significantly shorter PFS than those with low sCD155 levels (Figure 1C, D; Figure S1C, D). By contrast, there was no difference in PFS across sCD155 levels in patients with PDL-1 TPS less than 1% (Figure 1E). These findings indicate that elevated circulating sCD155 acts as a critical determinant of resistance to PD-1 blockade, particularly in tumors otherwise predicted to respond based on PD-L1 TPS. Collectively, these clinical data identify sCD155 as a systemic negative regulator of immunotherapy efficacy in NSCLC.

**Figure 1.**
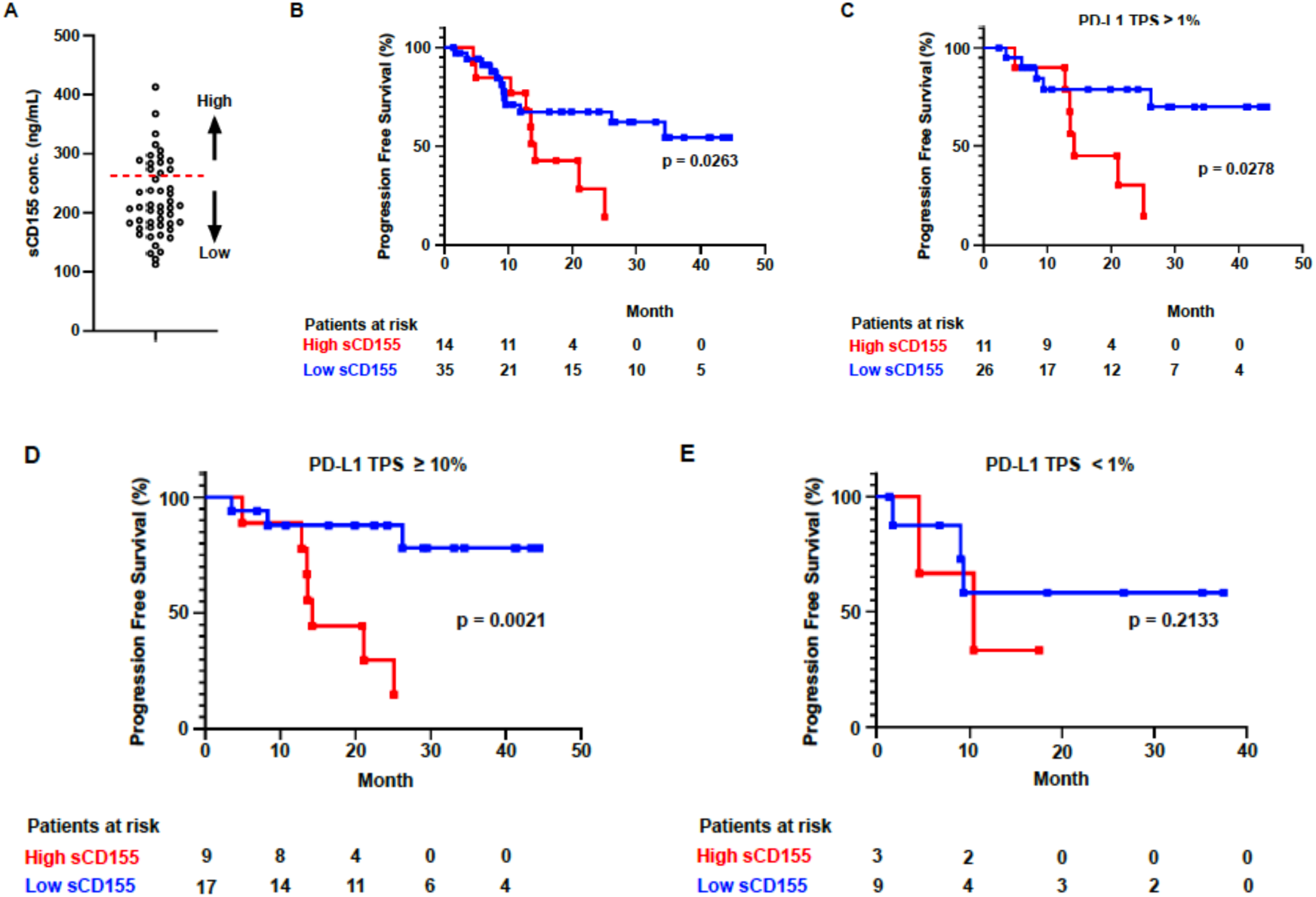
Elevated plasma sCD155 correlates with poor prognosis and anti-PD-1 resistance in NSCLC patients. **(A)** Pretreatment plasma sCD155 concentrations in patients with NSCLC (n = 49) receiving pembrolizumab monotherapy or combination therapy. Patients were stratified into high (≥260 ng/mL, n = 14) and low (<260 ng/mL, n = 35) sCD155 groups. **(B–E)** Kaplan-Meier curves for progression-free survival (PFS) comparing high and low sCD155 groups in the total cohort (B), patients with PD-L1 TPS ≥ 1% (C), ≥ 10% (D), and <1% (E). Statistical significance was determined by the log-rank test.

### sCD155 serves as a soluble immune checkpoint driving immunotherapy resistance

To mechanistically define the role of sCD155, we engineered murine tumor cell lines (B16/BL6 and MC38) to ectopically secrete the chimeric sCD155 consisting of the extracellular portion of mouse CD155 fused with the intracellular portion of human CD155 and the CT26 tumor cell line secreting only the extracellular portion of mouse CD155, as endogenous murine CD155 lacks soluble isoforms. Expression of sCD155 did not alter surface levels of membrane-bound CD155, PD-L1, or PD-L2 (Figure S2A), nor did it affect tumor growth in immunodeficient hosts, excluding tumor-intrinsic effects (Figure S2B-D). In contrast, in immunocompetent mice, sCD155-producing tumors exhibited markedly accelerated progression compared with control tumors (Figure 2A), demonstrating that sCD155 suppresses antitumor immunity in vivo. The circulating sCD155 level increased in proportion to tumor burden (Figure S2E, F), consistent with a systemic mode of action.

**Figure 2.**
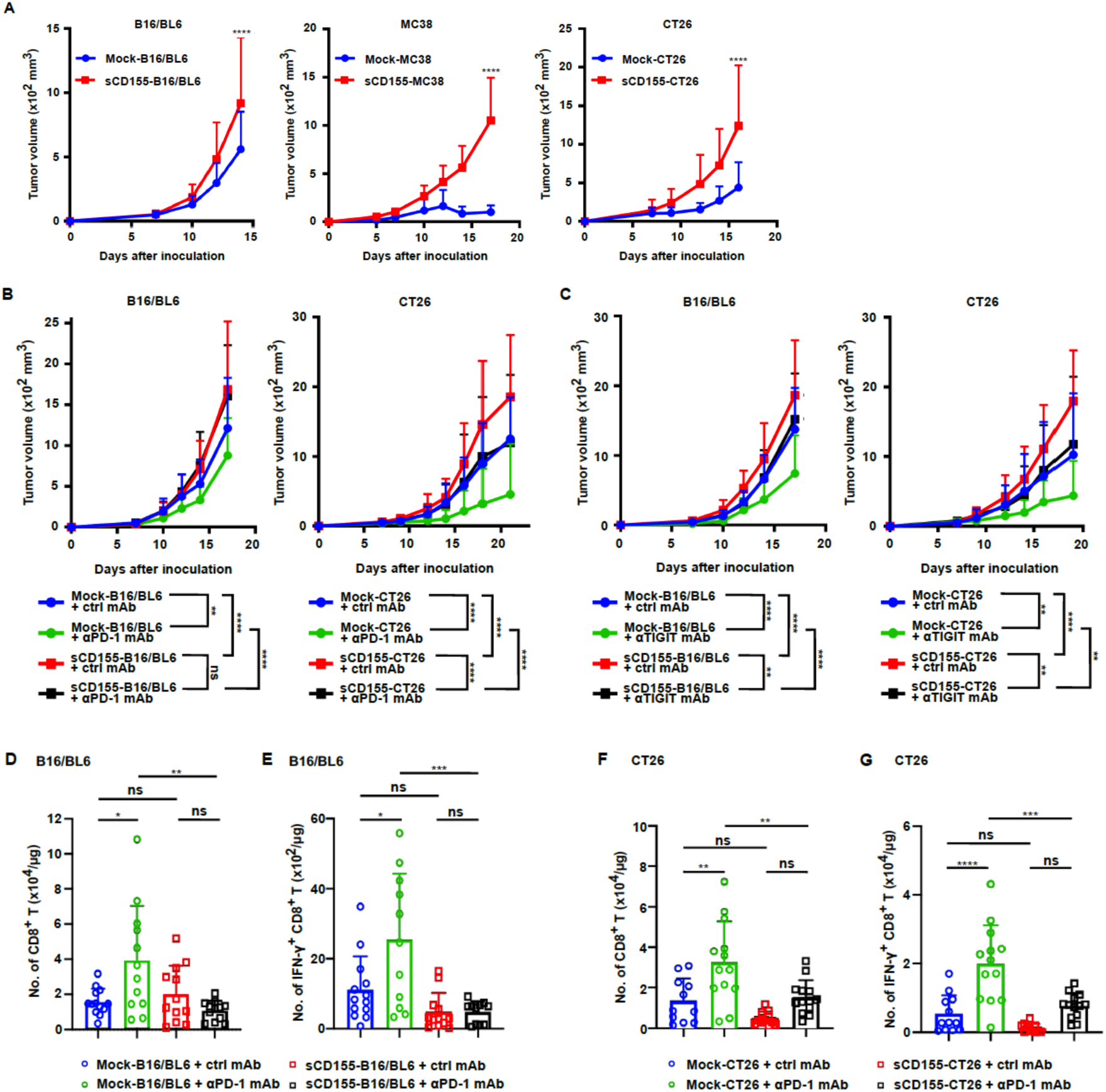
sCD155 functions as a soluble immune checkpoint to drive immunotherapy resistance. Cumulative tumor growth curves of mock-transfected or sCD155-producing B16/BL6 (mock, n = 15; sCD155, n= 17), CT26 (mock, n = 13; sCD155, n = 16), and MC38 (mock, n = 8; sCD155, n= 11) tumors in syngeneic WT mice. **(B and C)** Growth of mock or sCD155-producing B16/BL6 (n = 20 or 21 (B); n =18 or 19 (C)) and CT26 (n = 20 or 21 (B); n =14 - 16 (C))) tumors treated with control IgG, anti-PD-1 mAb (B), or anti-TIGIT mAb (C). **(D-G)** Numbers of tumor-infiltrating CD8^+^ T (D and F) and IFN-γ^+^ CD8 T (E and G) cells in mock and sCD155-producing B16/BL6 (D, E) and CT26 (F, G) in mice treated with control mAb (mock (n = 13 (D and E) and 11 (F and G); sCD155 (n = 13 (D), 11 (E), and 13 (F, G)) or anti-PD-1 mAb (mock (n = 13 (D, F, and G) and 12 (E); sCD155 (n = 13 (D, F and G) and 11 (E)). The data were pooled from two or three (A), four (B), three (C), and two (D-G) independent experiments. Statistical analyses were performed by two-way ANOVA (A-C) or one-way ANOVA (D, E). Error bars indicate mean ± SD. ns, not significant; *p < 0.05; **p < 0.01; ****p < 0.0001.

Strikingly, therapeutic blockade of PD-1 effectively controlled mock tumors but failed to restrain sCD155-expressing tumors across multiple tumor models (Figure 2B). Similarly, sCD155 diminished the antitumor efficacy of TIGIT blockade (Figure 2C). Analysis of tumor-infiltrating lymphocytes in the B16/BL6 tumor revealed that sCD155 abrogated the expansion and activation of cytotoxic T cells induced by PD-1 blockade, as evidenced by reduced IFN-γ, whereas NK cells and regulatory T cells were not significantly altered (Figures S2G-J).

Collectively, these findings demonstrate that sCD155 suppresses effector lymphocyte activation and confers resistance to immune checkpoint blockade. Unlike conventional membrane-bound checkpoints that require cell–cell contact, sCD155 acts in a soluble and systemic manner, thereby extending checkpoint regulation beyond local interactions. These data establish sCD155 as a soluble immune checkpoint that drives resistance to ICI therapy.

### sCD155 rewires checkpoint signaling by disabling DNAM-1–dependent activation

To identify the receptor pathway responsible for sCD155-mediated immune suppression, we evaluated tumor progression in mice deficient for CD155-binding receptors. The growth advantage of B16/BL6 and CT26 tumors conferred by sCD155 was completely abolished in DNAM-1–deficient mice, whereas TIGIT- and CD96-deficient hosts remained susceptible to sCD155-mediated B16/BL6 progression (Figures 3A-D). Depletion of CD8⁺ T cells or NK cells further demonstrated that either CD8⁺ T cells, NK cell, or both are required for sCD155-dependent progression of B16/BL6 and CT26 tumors (Figures 3E-H), identifying DNAM-1–expressing cytotoxic lymphocytes as primary targets. These results establish DNAM-1 as the critical molecular effector through which sCD155 exerts its immunosuppressive activity.

**Figure 3.**
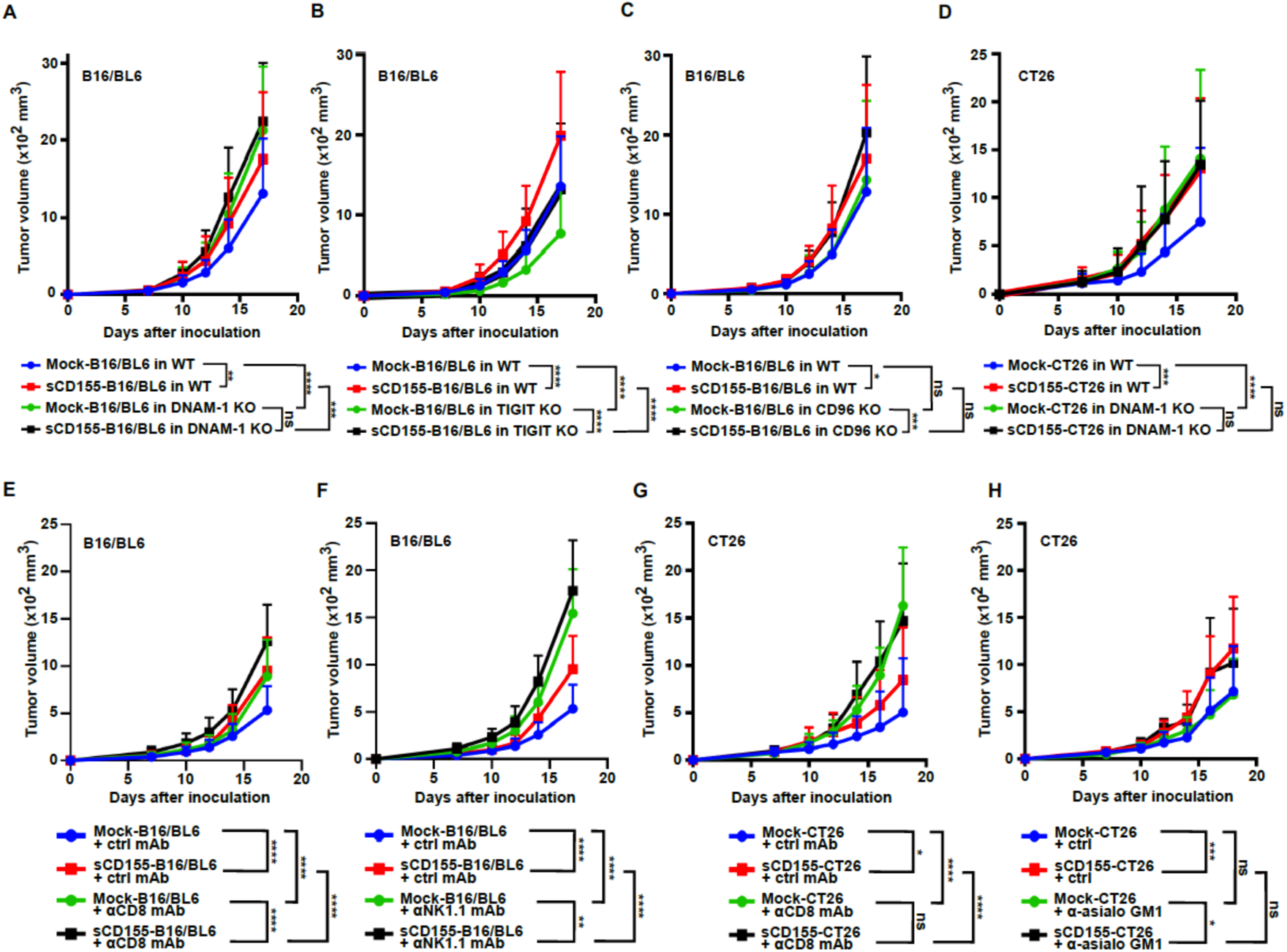
sCD155-mediated immune suppression is strictly dependent on the DNAM-1 signaling axis. **(A-D)** Growth kinetics of mock or sCD155-producing B16/BL6 (A-C) or CT26 (D) tumors in WT (mock-B16/BL6 (n = 25 (A), 23 (B), and 19 (C)) and mock-CT26 (17 (D)); sCD155-B16/BL6 (n = 23 (A), 22 (B), and 19 (C) and sCD155-sCD155-CT26 (17 (D)), *Cd226* (DNAM-1) KO (mock-B16/BL6 (n = 19 (A) and mock-CT26 (15 (D)); sCD155-B16/BL6 (19 (A)) and sCD155-CT26 (14) (D)), *Tigit* KO (mock B16/BL6 (n = 19) and sCD155-B16/BL6 (19)) (B) and *Cd96* KO (mock-B16/BL6 (n = 17); sCD155-B16/BL6 (16)) (C). **(E-H)** Tumor growth curves of B16/BL6 or CT26 tumors in mice treated with control mAb (mock-B16/BL6 (n = 14 (E and F)) and mock-CT26 13 (G and H)); sCD155-B16/BL6 (n = 14 (E and F)) and sCD155-CT26 (13 (G and H)), anti-CD8 mAb (mock-B16/BL6 (n = 13) (E) and mock-CT26 (13) (G); sCD155-B16/BL6 (n = 14) (E) and sCD155-CT26 (13) (G)), or anti-NK1.1 mAb (moc-B16/BL6 and sCD155-B16/BL6 (n =14) (F) and anti-asialo GM1 (mock-CT26 (n =10); sCD155-CT26 (9)) (G). Data are pooled from three (A, B, and D) and two (C and E-H) independent experiments. Statistical analyses were performed by two-way ANOVA. Error bars indicate means ± SD. ns, not significant; *P < 0.05; **P < 0.01; ***P < 0.001; ****P <0.0001

Given the central role of DNAM-1 as an activating receptor functionally opposed by PD-1 and TIGIT pathways, these findings indicate that sCD155 acts by selectively impairing DNAM-1–mediated signaling. Thus, sCD155 rewires checkpoint signaling and converts otherwise responsive tumors into an immune-refractory state by functionally uncoupling activating pathways downstream of ICI therapy.

### Targeting sCD155 by antibody restores cytotoxic lymphocyte-mediated antitumor immunity across species

To therapeutically target sCD155, we generated a monoclonal antibody (anti-sCD155 mAb) that selectively recognizes the intracellular portion of the human CD155 within the chimeric sCD155 without binding membrane-bound CD155 (Figure 4A) (29). Administration of anti-sCD155 mAb robustly suppressed the growth of sCD155-producing tumors but had no effect on control tumors lacking sCD155 (Figure 4B-D, Figures S3A, B), demonstrating high on-target activity. Ex vivo analysis demonstrated that treatment with this mAb markedly enhanced effector functions of both CD8⁺ T cells and NK cells within tumors, increasing cytokine production and degranulation activity (Figure S3C). Consistent with this, the depletion of CD8⁺ T cells and NK cells confirmed that both populations are essential for the therapeutic efficacy of the anti-sCD155 mAb (Figure 4B-D).

**Figure 4.**
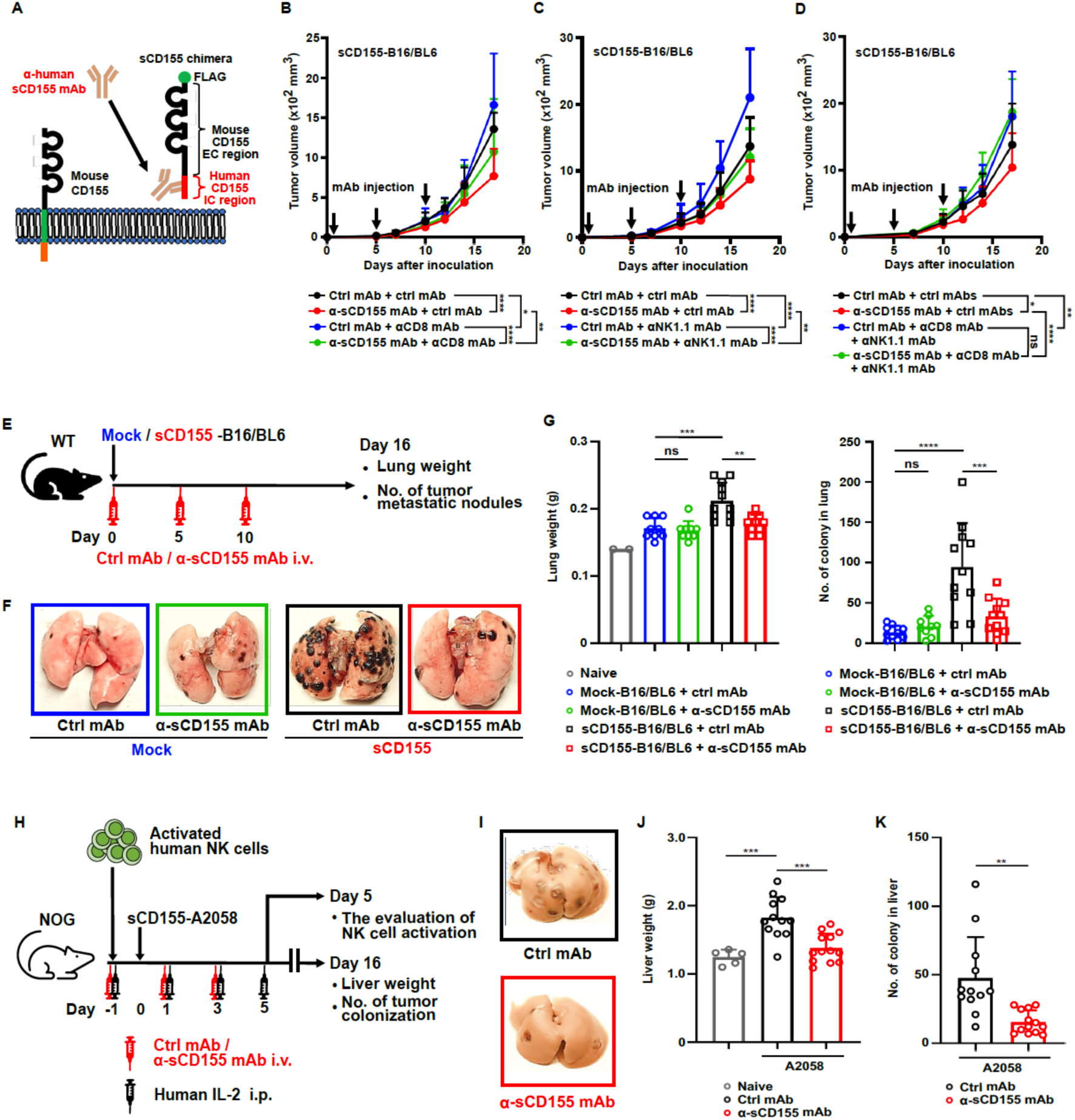
Selective targeting of sCD155 restores antitumor immunity and inhibits metastatic colonization. **(A)** Schematic representation of the anti-sCD155 mAb targeting the human CD155 intracellular (IC) domain of the chimeric sCD155 protein expressed by murine tumor cells. **(B)** Growth of sCD155-producing B16/BL6 tumors in mice treated with control mAb plus either control mAb (B-D) (n = 11 (B), 12 (C), and 10 (D), anti-CD8 mAb (B) (n =12), anti-NK1.1 mAb (C) (n =12), or both (D), or anti-sCD155 mAb plus either control mAb (B-D) (n = 12 (B), 11 (C), and 10 (D), anti-CD8 mAb (B) (n = 12), anti-NK1.1 (C) (n = 12)or both (D) (n = 10). Treatment was administered on Days 0, 5, and 10. **(E–G)** Schematic representation of the lung tumor colonization model using mock and sCD155-producing B16/BL6 cells (E). Representative lung images (F) and quantification of metastatic foci (G) (n = 9 or 10 per group). **(H and I)** Schematic representation of liver metastasis model using sCD155-producing A2058 tumor in humanized mice treated with contril mAb or anti0sCD155 mAb (H). Representative liver images (I) and quantification of liver weight and metastatic foci (J) (naïve (n = 5) (J); control mAb (10) (J and K); anti-sCD155 mAb (13) (J and K)). Data are pooled from two (B-D and G) and four (J) independent experiments. Error bars indicate means ± SD. Statistical analyses were performed by two-way ANOVA (B-D) and by one-way ANOVA (G and J). ns, not significant; *P < 0.05; **P < 0.01; ***P < 0.001; ****P < 0.0001

In an NK cell–dependent lung colonization model, sCD155 expression promoted metastatic colonization, whereas anti-sCD155 mAb significantly reduced tumor burden (Figures 4E-G). Importantly, the antitumor activity of anti-sCD155 mAb was fully recapitulated in humanized mouse models reconstituted with activated human NK cells following the injection of sCD155-producing human melanoma A2058 (Figure S3D). In this model, treatment increased accumulation and activation of human NK cells and suppressed metastatic progression (Figures 4H-K, Figures S3E-G). Collectively, these findings establish that targeting sCD155 restores antitumor immunity across murine and human immune systems.

### Antibody-mediated clustering of sCD155 converts immune suppression into DNAM-1 activation

Mechanistically, anti-sCD155 mAb did not deplete sCD155 from tumor interstitial fluid (Figure S4A), indicating a non-neutralizing mode of action. Anti-sCD155 mAb treatment failed to reduce tumor burden in DNAM-1–deficient settings, whereas its efficacy remained intact despite TIGIT deficiency (Figure 5A, B, Figure S4B, C). Instead, DNAM-1 reporter assays revealed that immune complexes formed between anti-sCD155 mAb and sCD155 triggered potent DNAM-1 signaling only when anti-sCD155 mAb was immobilized (Figure S4D). In contrast, while TIGIT signaling in TIGIT reporter cells was induced under plate-coated anti-sCD155 mAb and sCD155 conditions, the intensity of GFP expression was minimal compared with the positive control stimulation using anti-TIGIT mAb and CD155-Fc (Figure S4E). Furthermore, immobilized anti-sCD155 mAb enhanced cytokine production in both CD8^+^ T cells and NK cells (Figures S4F, G). Together, these results suggest that the sCD155 immune complex requires immobilization of the antibody’s Fc domain to activate DNAM-1 on cytotoxic lymphocytes.

**Figure 5.**
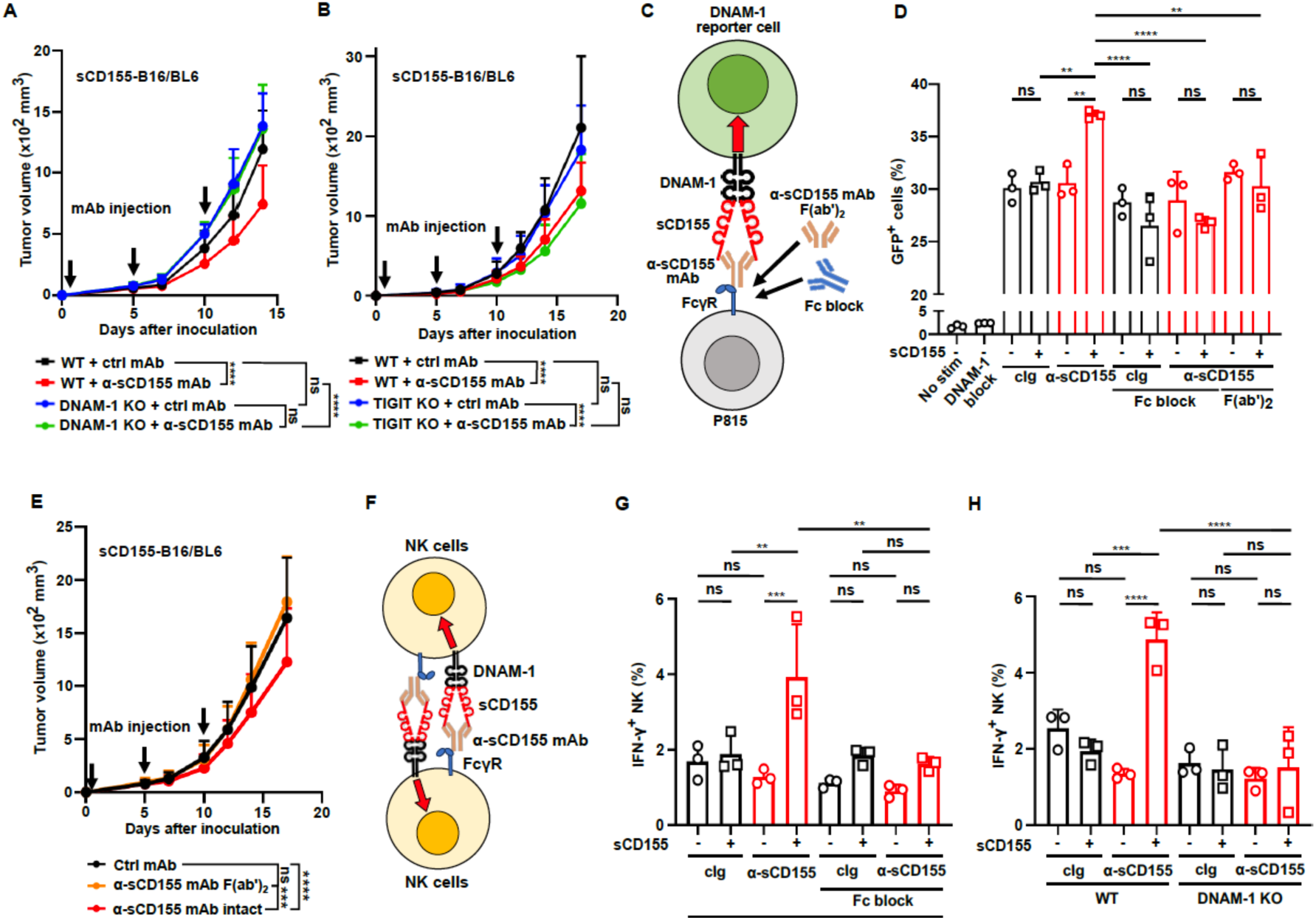
Antibody-mediated clustering of sCD155 via FcγR engagement converts immune suppression into DNAM-1 activation. **(A, B)** Cumulative tumor growth curves of sCD155-producing B16/BL6 tumors in WT, *Cd226* (DNAM-1) KO, and *Tigit* KO mice treated with control mAb (WT (n = 8 (A) and 11 (B)); DNAM-1 KO (8) (A); *Tigit* KO (13) (B)) or anti-sCD155 mAb (WT (n = 8) (A) and 9 (B); DNAM-1 KO (10); *Tigit* KO (10)). Treatment was administered on Days 0, 5, and 10. **(C and D)** Schematic representation (C) and quantitative analysis (D) of the DNAM-1 reporter cell co-culture assay with FcγR+ P815 cells. Reporter cells were stimulated with sCD155 or BSA in the presence of intact anti-sCD155 mAb, F(ab’)_2_ fragments, or Fc-block (n = 3 per group). (**E)** Growth of sCD155-producing B16/BL6 tumors in mice treated with intact anti-sCD155 mAb (n = 15), F(ab’)_2_ fragments (10), or control IgG (11). **(F-H)** Schematic and results of in vitro primary NK cell stimulation assays. NK cells from WT or DNAM-1 KO mice were stimulated with anti-sCD155 mAb via NK-NK interaction in the presence or absence of Fc-block (n = 3 per group). Data are pooled from two independent experiments (A, B, and E) and are representative of two independent experiments with similar results (D, G, and H). Statistical analyses were performed by two-way ANOVA (A, B, and E) or one-way ANOVA (D, G, and H). Error bars indicate means ± SD. ns, not significant; **p < 0.01; ***p < 0.001; ****p < 0.0001.

To test this hypothesis, we performed a coculture assay of DNAM-1 reporter cells with FcγR-expressing P815 cells. The results demonstrated a stringent requirement for FcγR engagement for DNAM-1 signaling, as *F(ab’*)*_2_* fragments or FcγR-blocking antibodies completely abolished DNAM-1 activation (Figures 5C, D). Consistently, treatment with *F(ab’)_2_* fragments of the anti-sCD155 mAb eliminated the antitumor efficacy in vivo (Figure 5E). Moreover, while cytokine production and cytolytic activity of primary NK cells―which abundantly express FcγR―were augmented when an sCD155 immune complex, this effect was abrogated by either FcγR blockade or DNAM-1 deficiency (Figure 5F-H). These findings demonstrate that antibody-mediated clustering of sCD155 via FcγR engagement converts a soluble inhibitory checkpoint into a potent DNAM-1–activating scaffold.

### sCD155 targeting synergizes with immune checkpoint blockade to convert resistance into durable tumor control

Given that sCD155 disables DNAM-1 signaling while anti-sCD155 mAb actively promotes its activation, we hypothesized that targeting sCD155 would potentiate the efficacy of existing ICIs. Indeed, combination therapy using the anti-sCD155 mAb alongside either PD-1 or TIGIT blockade produced significantly greater tumor suppression than any monotherapy alone (Figures 6A, B). Survival analyses further demonstrated that these combination treatments achieved the highest rates of durable tumor control (Figures 6C, D). Together, these data reveal that targeting the soluble checkpoint sCD155 rewires immune signaling to restore DNAM-1-dependent effector function, effectively converting immunotherapy resistance into therapeutic responsiveness.

**Figure 6.**
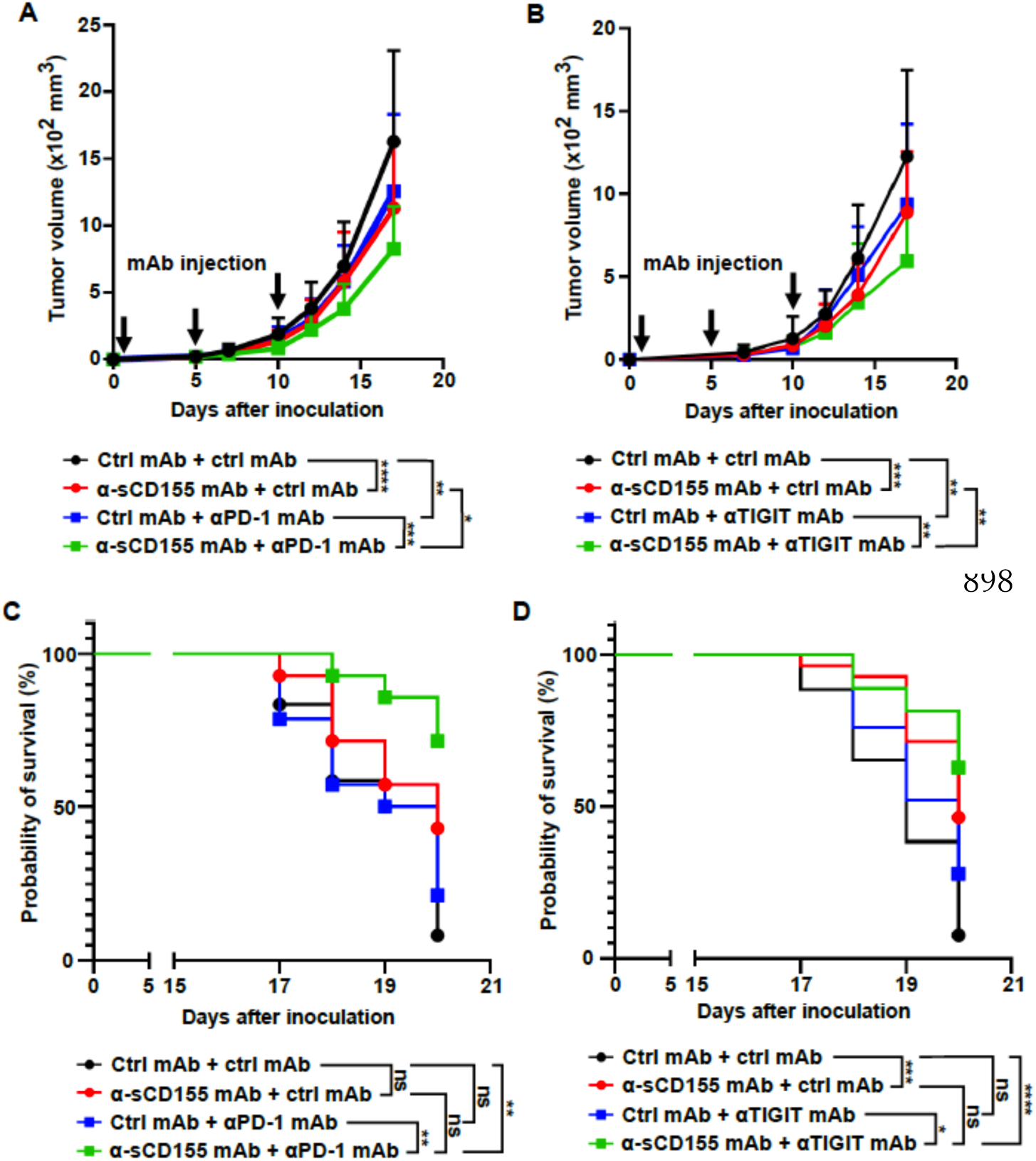
Targeting the sCD155–DNAM-1 axis synergizes with PD-1 and TIGIT blockade to achieve durable tumor control. **(A and B)** Cumulative tumor growth (A, B) and survival curves (C, D) of mice bearing sCD155-producing B16/BL6 tumors treated with combination therapy of anti-sCD155 mAb and either anti-PD-1 mAb (A, C) or anti-TIGIT mAb (B, D) (control mAb plus control mAb, n = 11 (A),12 (B and C), and 25 (D); control mAb plus anti-sCD155 mAb, n = 12 (A and B), 14 (C) and 25 (D); anti-PD-1 mAb plus control mAb, n = 11 (A) and 14 (C); anti-TIGIT mAb plus control mAb, n = 13 (B) and 28 (D); anti-PD-1 mAb plus anti-sCD155 mAb, n = 11 (A) and 14 (C); anti-TIGIT mAb plus anti-sCD155 mAb, n = 11 (B) and 27 (D)). Treatment was administered on Days 0, 5, and 10. Data are pooled from two (A-C) and four (D) independent experiments. Statistical analyses were performed by two-way ANOVA (A, B) and the log-rank test (C, D). Error bars indicate means ± SD. *p < 0.05; **p < 0.01; ***p < 0.001; ****p < 0.0001.

## Discussion

The CD155–DNAM-1 axis has traditionally been interpreted within a competitive receptor framework at the tumor cell surface, where elevated CD155 preferentially engages inhibitory receptors such as TIGIT and CD96, thereby suppressing cytotoxic lymphocyte function (11,30–32). While this model partially explains the paradoxical association between high CD155 expression and poor clinical outcomes despite the activating nature of DNAM-1 (33), it fails to account for the systemic immune dysfunction observed in cancer patients. In this study, we fundamentally revise this paradigm by identifying sCD155, whose levels in the circulation are positively associated with CD155 expression on tumors, as an active immunosuppressive ligand that directly disables DNAM-1 signaling, independent of inhibitory receptor competition. Moreover, we demonstrate that immune complex formation converts sCD155 into a potent immunoactivating scaffold, establishing ligand valency and spatial organization as decisive regulators of DNAM-1 signaling quality.

DNAM-1 is a central costimulatory receptor expressed on NK cells and CD8⁺ T cells that promotes immune synapse formation, cytotoxic granule polarization, and inflammatory cytokine production (14,15,16,34,35). Genetic ablation or functional disruption of DNAM-1 profoundly impairs antitumor immunity across multiple tumor models, underscoring its essential role in immune surveillance ‘18’. Yet, in human malignancies, CD155 is frequently overexpressed and correlates with immune dysfunction and unfavorable prognosis (8,10,36–40). Our findings reveal that beyond membrane-bound ligand competition, circulating sCD155 establishes a systemic suppressive layer that actively reprograms DNAM-1 signaling competence.

Mechanistically, we show that monomeric or low-valency sCD155 engages DNAM-1 in a non-productive manner, dispersing receptor organization and attenuating phosphorylation of key downstream signaling intermediates. This uncoupling of ligand binding from activating signal propagation effectively renders DNAM-1 functionally silent despite high receptor occupancy. Such behavior aligns with emerging principles in immunoreceptor systems where low-avidity or spatially diffuse engagement enforces tolerance, anergy, or exhaustion rather than activation (41,42). Thus, sCD155 functions not merely as a passive decoy but as an active suppressor that reshapes cytotoxic lymphocyte signaling landscapes.

In striking contrast, immune complex formation by anti-sCD155 mAb fundamentally alters the signaling outcome of sCD155 engagement. By enforcing multivalent DNAM-1 clustering, these immune complexes restore receptor microdomain assembly, sustain activating phosphorylation cascades, and reinvigorate NK and CD8⁺ T cell effector functions. This valency-dependent signaling switch parallels well-established paradigms in T cell receptor (TCR) and Fc receptor biology, where spatial organization and receptor density dictate qualitative signaling outputs (41,42). Our data extend these concepts to the DNAM-1 axis, positioning ligand architecture as a master regulator of immune activation versus suppression.

Clinically, elevated CD155 expression was prevalent across murine tumor models and human cancer patient cohorts, correlating strongly with impaired cytotoxic lymphocyte function. While CD155 has previously been reported as a biomarker associated with advanced disease and poor prognosis (39,40,43, 44), its mechanistic contribution remained undefined. Our work provides a compelling rationale for sCD155 as a therapeutic target-not merely to remove a systemic brake, but to actively fuel antitumor immunity.

Importantly, our findings also establish sCD155 as a key determinant of resistance to ICIs. Although checkpoint blockades release inhibitory receptor constraints, effective antitumor immunity ultimately requires intact activating receptor signaling. By functionally silencing DNAM-1 despite receptor engagement, elevated sCD155 creates a parallel suppressive axis that persists even in the presence of PD-1/PD-L1 or TIGIT blockade. This mechanism offers a direct explanation for why a substantial fraction of patients fail to respond to ICI despite apparent relief of inhibitory signaling pathways, consistent with studies showing that exhausted or activation-deficient lymphocyte states limit checkpoint therapy efficacy (45–47).

Strikingly, conversion of sCD155 into immune complexes not only restored DNAM-1 signaling but markedly enhanced responsiveness to checkpoint blockade in preclinical models. Through multivalent receptor engagement and the reconstitution of productive signaling microclusters, immune complexes effectively re-sensitized cytotoxic lymphocytes to ICI-driven reinvigoration. These findings position sCD155 not merely as a bystander biomarker of immune dysfunction, but as an active driver of immunotherapy resistance, while its immune complex form emerges as a potent determinant of therapeutic responsiveness.

A central conceptual advance of this study is the recognition that ligand architecture governs immune fate decisions within the CD155–DNAM-1 system. Monovalent soluble ligand enforces immune paralysis, whereas multivalent ligand organization restores activation. This dynamic signaling rheostat reframes the axis from a static receptor competition model into a spatially regulated signaling system capable of switching between suppression and activation. Such regulation may represent a general principle of immune tuning in chronic disease states where soluble immune ligands accumulate.

These findings carry profound implications for therapeutic strategies targeting the CD155 axis. While current approaches primarily focus on blocking inhibitory receptors such as TIGIT, our data indicate that restoration of activating receptor competence is equally essential for durable immune reinvigoration. Neutralization of sCD155 alone may be insufficient; rather, its enforced multimerization into activating complexes directly restores DNAM-1 function and synergizes with checkpoint blockade. Conversely, antibody formats that merely sequester soluble ligands without promoting productive clustering may fail to reverse immune suppression or could even perpetuate signaling dysfunction. Rational ligand-targeting strategies must therefore consider spatial signaling outcomes in addition to receptor occupancy.

Beyond cancer, soluble ligand–mediated modulation of receptor signaling via valency-dependent mechanisms may have broader relevance in chronic infection, autoimmunity, and inflammatory disorders, where elevated soluble immune ligands are common. Future work should explore whether analogous systems govern exhaustion and functional tuning across other activating receptor networks.

In conclusion, we uncover sCD155 as a previously unrecognized systemic immunosuppressive ligand that actively disables DNAM-1 signaling and promotes resistance to immune checkpoint therapy. Paradoxically, immune complex formation converts this same ligand into a potent immunoactivator that restores receptor clustering, reinvigorates cytotoxic lymphocyte function, and enhances responsiveness to immunotherapy. This work reframes the CD155–DNAM-1 axis as a dynamic spatial signaling rheostat governed by ligand valency and organization, revealing new therapeutic opportunities to reprogram antitumor immunity and overcome immunotherapy resistance.

## Limitations of the study

While we establish functional consequences of DNAM-1 spatial reorganization, high-resolution structural characterization of sCD155–DNAM-1 complexes will be necessary to define precise clustering geometries and stoichiometries. Moreover, prospective clinical studies are required to determine whether circulating sCD155 predicts immunotherapy resistance or response to ligand-modulating strategies. Finally, the exact mechanisms by which sCD155 differentially regulates activating versus inhibitory receptor engagement within the broader receptor network warrant further investigation.

## Acknowledgments

This work was supported by JSPS KAKENHI (21H04836 to A.S., 21K19469 and 16H05169 to K.S). The authors thank M. Kaneko for administrative support.

## Author Contributions

S.K., T.M., K.O, N.T., G.O., N.I., S.G., and E.S. performed experiments.

S.K., T.M., G.O., S. K., A. I-M., C.N-O., K.S., and A.S. performed data analysis and project planning.

H.T. provided an essential material.

S.K., T.M, K.S., and A.S. contributed to writing the manuscript.

K.S. and A.S. supervised the overall project.

## Materials and Methods

### Patients and Sample Collection

We retrospectively enrolled patients with advanced non-small cell lung cancer (NSCLC) who received either pembrolizumab monotherapy or combination therapy with chemotherapy as first-line treatment at the National Cancer Center Hospital East. Eligible patients met the following criteria:

- Performance Status: ECOG score of 0–2.
- Measurable Disease: Confirmed by RECIST version 1.1.
- Treatment Adherence: Receipt of at least two treatment cycles.

Plasma samples were collected from all participants immediately prior to the initiation of therapy. All patients provided written informed consent, and the study protocol was approved by the Institutional Research Ethics Committee in accordance with the Declaration of Helsinki and ICH-GCP guidelines.

### Clinical Protocols and Endpoints

Pembrolizumab was administered intravenously at 200 mg every 3 weeks or 400 mg every 6 weeks for monotherapy, and 200 mg every 3 weeks in combination regimens.

Chemotherapy dosing followed standard clinical practice:

- Carboplatin: AUC 5–6.
- Pemetrexed: 500 mg/m².
- Paclitaxel: 175–200 mg/m².

The primary endpoint was progression-free survival (PFS), assessed to evaluate the association with plasma sCD155 levels. Patients were followed longitudinally for survival and safety, including physical examinations, vital signs, and laboratory testing. Tumor PD-L1 expression was quantified using the Tumor Proportion Score (TPS), calculated by (PD-L1-positive tumor cells/all viable tumor cells) x 100.

### Mice

C57BL/6J and BALB/c wild-type mice were purchased from CLEA Japan (Tokyo, Japan). NOD.Cg-*Prkdc^scid^Il2rg^tm1Sug^*/ShiJic mice were obtained from the Central Institute for Experimental Animals (Kawasaki, Japan). *Cd226*-/-, *Tigit*-/-, and *Cd96*-/- mice were generated as previously described (24) and backcrossed onto the appropriate genetic backgrounds for at least ten generations. All mice were maintained under specific pathogen-free (SPF) conditions at the Laboratory Animal Resource Center of the University of Tsukuba. Mice were used for experiments at 8–12 weeks of age, with age- and sex-matched littermates used as controls where applicable. All animal experiments were performed in accordance with relevant institutional and national guidelines and were approved by the Institutional Animal Care and Use Committee (IACUC) of the University of Tsukuba (Approval No. 22-155 and 23-120).

### Cell lines and generation of sCD155-producing cells

To generate mouse sCD155-producing CT26 cells, a sequence combining the DYKDDDDK flag tag and exons 1–5 of mouse *Cd155* was ligated into the pMXs-IRES-GFP retroviral vector (CBL, RTV-013). This vector was transduced into the CT26 mouse colorectal cancer cell line using Polybrene infection/transfection reagent (#TR-1003, Sigma-Aldrich). As a control, an empty pMXs-IRES-GFP vector was transduced. Transduced cells were bulk-sorted using a FACSAria III (BD Biosciences) based on GFP expression levels. The resulting populations were termed “sCD155-CT26” and “mock-CT26,” respectively. To generate human sCD155-producing A2058 cells, the sequence of human *CD155ß* was ligated into the pMXs-IRES-GFP vector. To generate the “sCD155 chimera” consisting of the mouse CD155 extracellular region and human CD155 intracellular region, a sequence containing the DYKDDDDK flag, exons 1–5 of mouse *Cd155*, and exons 7–8 of human *CD155* was assembled using the NEBuilder HiFi DNA Assembly Cloning Kit (#E5520S, New England Biolabs). This vector was transduced into B16/BL6 and MC38 cells. The establishment of these sCD155-producing tumor lines followed the same procedure as for sCD155-CT26.

B16/BL6 cells were cultured in RPMI-1640 (#189-02025, Wako) supplemented with 10% fetal bovine serum (FBS; #FB-1061/500, Biosera), 10 mM HEPES (#15630-130, Gibco), 1 mM sodium pyruvate (#11360-097, Gibco), 1x MEM Non-Essential Amino Acids (#11140-076, Gibco), and 100 U/mL penicillin-streptomycin (#G1146-100ML, Sigma-Aldrich). CT26, MC38, and A2058 cells were cultured in Dulbecco’s Modified Eagle Medium (DMEM; #044-29765, Wako) supplemented with the same components as the RPMI-1640 medium. Cells were maintained in a humidified incubator at 37°C with 5% (for B16/BL6) or 10% (for CT26, MC38, and A2058). All cell lines were regularly tested for *Mycoplasma* using the MycoAlert *Mycoplasma* Detection Kit (#LT07-418, Lonza) to ensure they remained contamination-free.

NFAT-GFP reporter Ba/F3 cells were kindly provided by Dr. Hisashi Arase (Osaka University, Osaka, Japan). Mouse DNAM-1 and TIGIT reporter cells were generated by retroviral transduction of mouse DNAM-1-FcRγ and TIGIT-FcRγ chimeric proteins into the reporter cell line, as previously described (48). These lines were maintained in RPMI-1640 supplemented with 10% WEHI-3 cell culture supernatant and 10% FBS.

### Analysis of CD155 and sCD155 Isoform Expression

To investigate the correlation between membrane-bound CD155 and its soluble isoforms, we reanalyzed mRNA expression profiles using a previously described dataset (26). Specifically, the relative expression of (encoding CD155) and its soluble variants (encoding sCD155) was compared between primary tumor tissues and adjacent healthy tissues. This analysis included data from 16 patients across multiple cancer types, including colorectal, gastric, and breast cancers.

### Sandwich ELISA for sCD155 in cancer patients

Plasma samples from patients with NSCLC were collected immediately prior to the initiation of immune checkpoint inhibitor therapy. Ninety-six–well plates were coated overnight with 20 ng/well of anti-CD155 antibody in carbonate-bicarbonate buffer at 4°C. After washing and blocking, plasma samples were incubated for 1 h at room temperature (RT). Bound sCD155 was detected using a rabbit anti-PVR antibody, followed by a biotinylated secondary antibody and streptavidin-HRP. The colorimetric signal was developed using TMB substrate, neutralized with sulfuric acid, and the absorbance was measured at 450 nm using a microplate reader.

### Sandwich ELISA for sCD155 measurement in mouse serum

Serum samples were obtained via centrifugation of whole blood. For the ELISA, 96-well plates were coated overnight with 200 ng/well of anti-mouse CD155 mAb (clone TX56), as previously described (24). After blocking, 100 µL of mouse serum samples or recombinant sCD155 chimera protein (for the standard curve) were added and incubated at room RT for 2 h. To detect sCD155 bound to TX56, 20 ng/well of rabbit anti-mouse CD155 polyclonal antibody (generated in-house) and HRP-conjugated donkey anti-rabbit IgG (1:10,000 dilution; #406401, BioLegend) were incubated for 1 h at RT. The reaction was developed with TMB substrate (#555214, BD), stopped with 0.5 M sulfuric acid (#28-5940-5-500ML, Sigma-Aldrich), and the absorbance was measured at 450 nm.

### Generation of sCD155-producing cell lines

Mouse and human sCD155 constructs were cloned into the pMXs-IRES-GFP retroviral vector and transduced into CT26, B16/BL6, MC38, and A2058 cells. Successfully transduced populations were sorted based on GFP expression using a FACSAria (BD Biosciences).

### Generation of F(ab’)2 fragments of anti-sCD155 mAb

To prepare *F(ab’)_2_* fragments, the anti-sCD155 mAb (clone 29) was enzymatically digested using the Pierce Mouse IgG1 Fab and *F(ab’)_2_* Preparation Kit (catalog #44980, Thermo Fisher Scientific) according to the manufacturer’s instructions. The resulting fragments were purified to remove intact IgG and Fc fragments, and their purity was confirmed by SDS-PAGE under non-reducing and reducing conditions.

### In vivo mouse subcutaneous tumor models

To establish subcutaneous tumor models, CT26 cells (5 x 10^6^ cells in 200 µL PBS), either mock-transfected or sCD155-producing, were subcutaneously inoculated into the flanks of male BALB/c or NOG mice. Similarly, B16/BL6 cells (5 x 10^5^ cells) or MC38 cells (1 x 10^6^ cells), expressing either mock or sCD155 chimera, were injected into male C57BL/6J or NOG mice.

For immune cell depletion, mice received weekly intraperitoneal (i.p.) injections starting from day −1 relative to tumor inoculation: 100 µg of rabbit anti-asialo GM1 (Fujifilm, #014-09801) for BALB/c mice, 100 µg of anti-mouse NK1.1 Ab (clone PK136, generated in-house), or 100 µg of anti-mouse CD8 Ab (clone 53–6.7, generated in-house).

For ICI treatment, 200 µg of anti-mouse PD-1 (clone RMP1-14, #BE0146, Bio X Cell) or anti-TIGIT Ab (clone TX99 [49]) was administered i.p. twice weekly for the B16/BL6 model and once weekly for the CT26 model, starting from day 0. Intact anti-sCD155 mAb or its *F(ab’)_2_* fragment (100 µg) was administered intratumorally on days 0, 5, and 10. For combination therapy, 100 µg of anti-PD-1 Ab (i.p.) and 100 µg of anti-sCD155 mAb (intratumorally) were administered on days 0, 5, and 10. Tumor volume was measured three times per week, and mice were euthanized when tumor volumes exceeded 2,000 mm^3^.

### In vivo mouse lung colonization models

B16/BL6 cells (2 x 10^5^ cells in 100 µL PBS), either mock-transfected or producing sCD155 chimera, were intravenously inoculated into male C57BL/6J WT mice. Simultaneously, 200 µg of anti-sCD155 mAb or an isotype control mAb was intravenously administered. Subsequently, additional doses were administered on days 5 and 10. After 16 days, lungs were resected and washed with PBS. Fixed lungs were divided into lobes, and the number of tumor nodules was counted bilaterally.

### In vivo liver tumor colonization in a humanized mouse model

Human PBMCs were isolated from 60 mL of blood from healthy donors using Lymphoprep™ density gradient medium (#18061, STEMCELL Technologies). Human CD56+ NK cells were positively selected using CD56 MicroBeads and LS Columns (#130-050-401, Miltenyi Biotec). Isolated NK cells were activated with 300 U/1 x10^6^ cells of recombinant human IL-2 (#554603, BD) for 24 h. These activated NK cells (5 x 10^5^ cells/mouse) were then adoptively transferred into NOG mice. On day −1, 75,000 U of recombinant human IL-2 (#200-02, Peprotech) and 100 µg of isotype or anti-sCD155 mAb were administered. Every other day until day 3 (for IL-2) and day 5 (for mAbs), treatments were continued intravenously. On day 0, 1 x 10^6^ sCD155-producing A2058 cells were inoculated intravenously. On day 16, liver weight and the number of tumor colonies were evaluated. This study was approved by the ethical review board of the University of Tsukuba (No. 240221).

### Tumor dissociation and cell isolation

Tumor tissues were excised and minced into small pieces. The tissues were then incubated in RPMI-1640 medium supplemented with 10% FBS, 100 µg/mL collagenase from *Clostridium histolyticum* (#5138, Sigma-Aldrich), and 100 µg /mL DNase I (#LS002139, Worthington Biochemical) at 37°C for 30 min, followed by mechanical dissociation using a gentleMACS™ Dissociator (#130-093-235, Miltenyi Biotec) according to the manufacturer’s instructions. The resulting single-cell suspension was washed twice with complete RPMI medium and filtered through a 70 µm cell strainer.

For functional assays, isolated cells were incubated in complete RPMI containing 1,500-fold diluted GolgiStop™ (#554724, BD) and 100-fold diluted anti-mouse CD107a-BV711 (clone 1D4B, #121631, BioLegend) for 5 h. After staining dead cells with Zombie NIR™ Fixable Viability Kit (#423105, BioLegend) and performing Fc blocking, the cell suspension was stained with the following anti-mouse antibodies: NK1.1-FITC (clone PK136, #108706, BioLegend), CD25-PE (clone PC61.5, #50-0251, TOMBO biosciences), TCR-beta chain-PE/Cy7 (clone H57-597, #109222, BioLegend), Alexa Fluor® 700-CD8a (clone 53-6.7, #557959, BD), BV421™-CD45.2 (clone 104, #109832, BioLegend), and BV510™-CD4 (clone RM4-5, #100559, BioLegend).

For intracellular staining, the eBioscience™ Fixation/Permeabilization Concentrate (#00-5123-43, Thermo Fisher) and Diluent were used for Foxp3, while Cytofix/Cytoperm™ (#554722, BD) was used for cytokine detection. Cells were stained with Alexa Fluor® 488-FOXP3 (clone 150D, #320012, BioLegend) for Treg identification, and APC- or PE-conjugated anti-IFN-γ (clone XMG1.2) and anti-TNF (clone MP6-XT22) for cytokine profiling.

### Analysis of liver-infiltrating human NK cells

Liver tissues with tumor nodules were collected from NOG mice 5 days after tumor inoculation and homogenized using a glass homogenizer (#81-0781, SANSYO). The homogenate was diluted with 40% Percoll™ PLUS solution (#17544501, Cytiva) and overlaid onto 60% Percoll™ PLUS. Following density gradient centrifugation, the buffy coat containing mononuclear cells was collected, and red blood cells were lysed using ACK lysis buffer.

The resulting lymphocytes were incubated in complete RPMI supplemented with 2,000-fold diluted GolgiStop™ and 100-fold diluted anti-human CD107a-APC (clone H4A3, #328620, BioLegend) for 5 h at 37°C in a 5% CO_2_ incubator. Zombie NIR™ was used to exclude dead cells. For human NK cell surface staining, anti-human CD3-PE (clone HIT3a, #555340, BD), anti-human CD45-biotin (clone HI30, #555481, BD), SA-PE-Cy7 (#557598, BD), and anti-human CD56-V450 (clone B159, #560360, BD) were used. For fixation and permeabilization, Cytofix/Cytoperm™ was used according to the manufacturer’s instructions. The number and function of the infiltrating NK cells were then analyzed by flow cytometry.

### Reporter assay for DNAM-1 and TIGIT signaling

For the plate-bound stimulation assay, 96-well plates were coated overnight with 5 µg /mL of each antibody: anti-DNAM-1 (clone TX42, [50]), anti-TIGIT (clone TX99, [49]), anti-sCD155 (29), or an isotype control. Mouse DNAM-1 and TIGIT reporter cells (48) were pre-incubated on ice for 30 min with either 5 µg of BSA or recombinant sCD155 chimera protein per 1 ×10^5 cells. For soluble antibody stimulation, reporter cells were incubated with anti-sCD155 mAb or isotype control, followed by BSA or sCD155 treatment, respectively. After washing, 2 x 10^4^ reporter cells per well were cultured for 24 h at 37°C in a 5% CO_2_ incubator. Cells were then resuspended in PBS containing 3,000-fold diluted propidium iodide (PI; #P4864, Sigma-Aldrich) and analyzed by flow cytometry (FACS).

For the coculture assay, 3 µg of anti-sCD155 or control Ab was added to Fc γ R-expressing P815 mastocytoma cells (1 × 10^5^ cells). After washing away unbound antibodies, P815 cells were incubated on ice for 30 min with 5 µg of BSA or recombinant sCD155 chimera per 1 × 10^5^ cells. To investigate concentration dependency, 0 to 5 µg of BSA or recombinant sCD155 chimera was applied. Both P815 cells and reporter cells (2 × 10^4^ cells each) were cocultured for 24 h. Following incubation, P815 cells were identified by staining with CD117-APC (clone 2B8, #105812, BioLegend), which does not bind to DNAM-1 reporter cells. Finally, cells were resuspended in PBS containing PI and analyzed by FACS.

### In vitro stimulation of CD8^+^ T cells and NK cells

For the stimulation of primary CD8^+^ T cells, 96-well plates were co-coated overnight with 500 ng/well of anti-sCD155 mAb or an isotype control, alongside indicated concentrations (0 to 30 ng/well) of anti-CD3e mAb (clone 145-2C11, #100340, BioLegend). Primary CD8^+^ T cells were isolated from the spleens of male C57BL/6J WT mice and purified using CD8a (Ly-2) MicroBeads (#130-117-044, Miltenyi Biotec) and LS columns (#130-042-401, Miltenyi Biotec) according to the manufacturer’s instructions.

To stimulate primary NK cells, splenic NK cells were isolated by negative selection and activated according to an established protocol (24). Plates were coated overnight with 0 to 800 ng/well of anti-sCD155 mAb or an isotype control.

For functional assessment, 5 × 10^4^ treated CD8^+^ T cells or 2 × 10^4^ NK cells were seeded per well in complete RPMI supplemented with 2,000-fold diluted GolgiStop™ (#554724, BD) and 100-fold diluted anti-mouse CD107a-BV711 (clone 1D4B, #121631, BioLegend) for 5 h. Intracellular cytokine production (IFN-gamma and TNF) was analyzed by flow cytometry, as described in the TIL analysis workflow above.

### Flow cytometry analysis

Cell suspensions were pre-incubated with Fc Block (anti-mouse CD16/32, clone 2.4G2, BD Biosciences) for 10 min at 4°C to minimize non-specific binding prior to staining with indicated fluorochrome-conjugated mAbs or isotype-matched controls. In all experiments, doublets and dead cells were rigorously excluded using FSC-A/FSC-H gating and viability dye staining, respectively. Data were acquired on an LSRFortessa or FACSAria III flow cytometer (BD Biosciences) using BD FACSDiva Software v8. All data were analyzed using FlowJo v10 (FlowJo, LLC).

## Quantification and statistical analysis

Statistical analyses were performed using GraphPad Prism 9 (GraphPad Software, San Diego, CA, USA). Data are presented as mean ± SD or mean ± SEM as indicated in the figure legends. For comparisons between two groups, a two-tailed Student’s t-test was used. For multiple group comparisons, one-way or two-way ANOVA followed by appropriate post-hoc tests (e.g., Tukey’s or Sidak’s) was applied. Fisher’s exact test was used for categorical data where appropriate. A P-value < 0.05 was considered statistically significant.

---

**Supplemental Figure 1.**
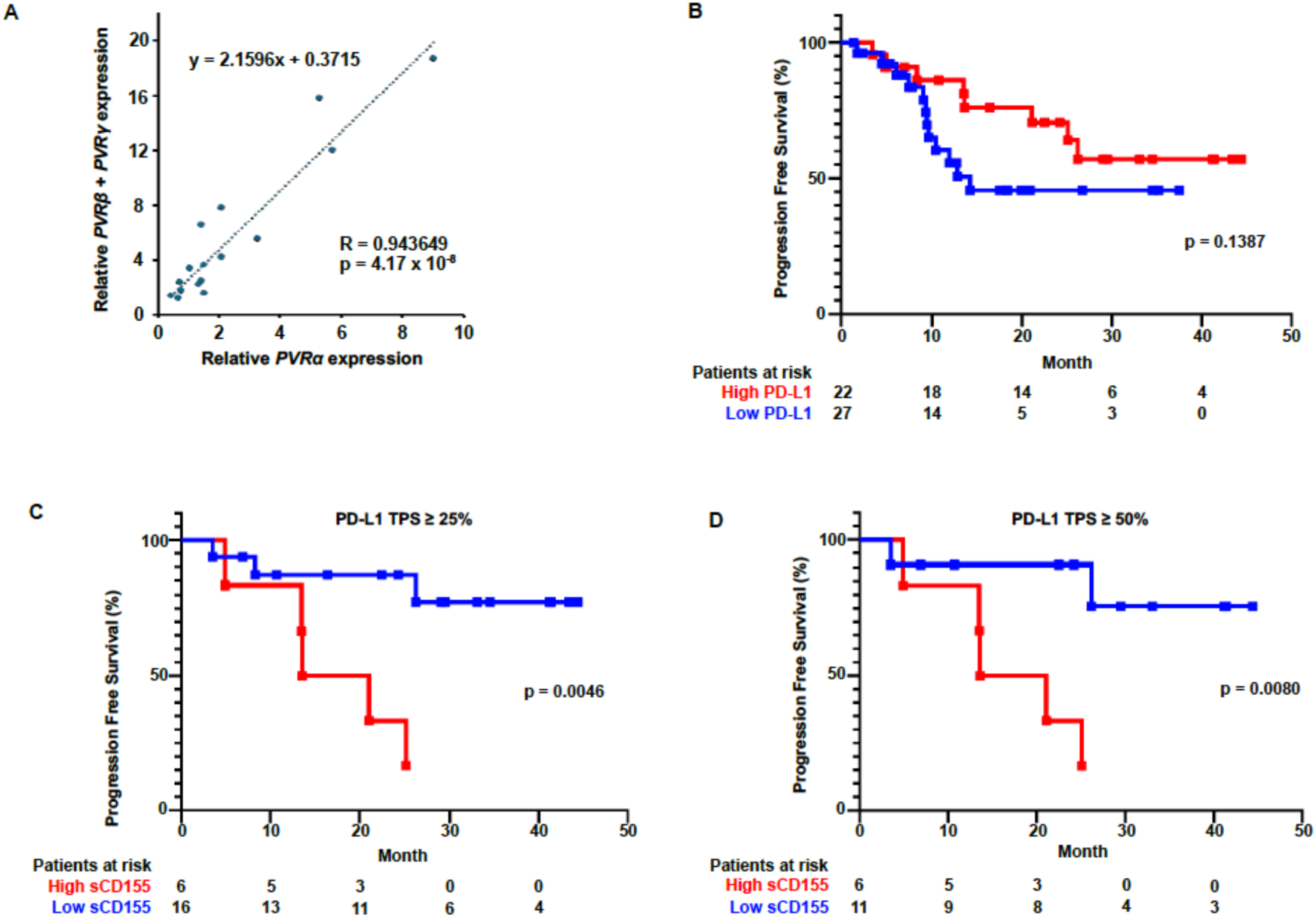
sCD155 levels and PD-L1 expression as determinants of PFS in NSCLC patients. **(A)** Association between the relative expression of CD155 (*PVRα*) and sCD155 (*PVRβ* plus *PVRγ*) mRNA in colorectal, gastric, and breast cancer compared with adjacent healthy tissues (n = 16). **(B)** Progression-free survival (PFS) of NSCLC patients stratified by high (≥ 50%) and low (<50%) PD-L1 tumor proportion score (TPS). **(C and D)** PFS analysis of patients with PD-L1 TPS with ≥ 25% (B) and ≥ 50% (C), further stratified by high and low sCD155 concentrations. Statistical significance was determined by the log-rank test (B-D).

**Supplemental Figure 2.**
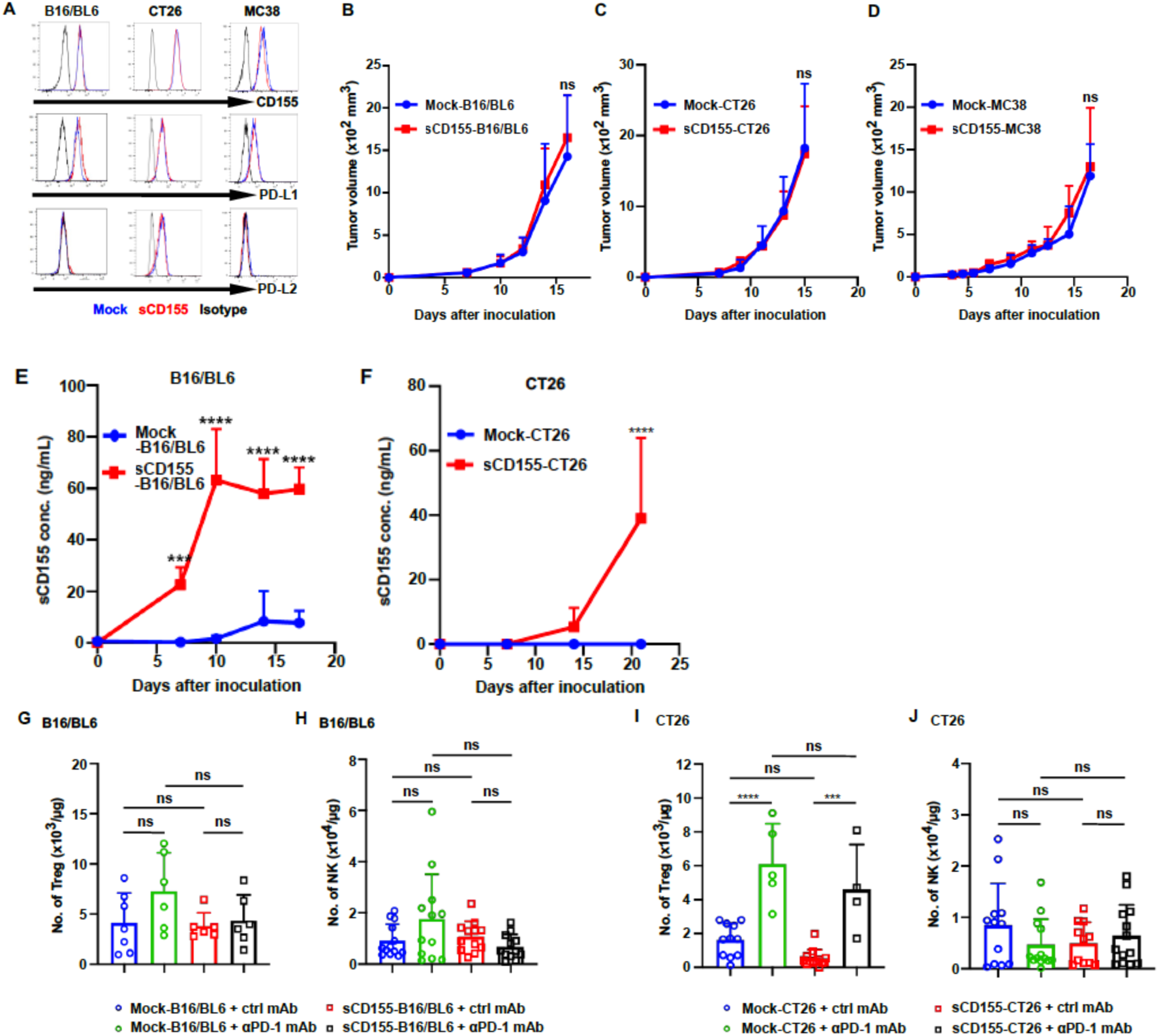
Characterization of sCD155-producing tumor models and systemic sCD155 accumulation. Flow cytometry analysis of membrane-bound CD155, PD-L1, and PD-L2 on mock and sCD155-expressing B16/BL6, CT26, and MC38 tumor cells. **(B-D)** Growth of mock (n = 13 (B), 3 (C), and 4 (D), sCD155-producing (n = 12 (B), 3 (C), and 4 (D) B16/BL6, CT26, and MC38 tumors in immunodeficient NOG mice (n = 3–4 per group). **(E and F)** Serum sCD155 concentrations in mice bearing mock or sCD155-expressing B16/BL6 and CT26 tumors (n = 3 or 4 per group). **(G-J)** Numbers of tumor-infiltrating Treg cells (G and I) and NK cells (H and J) in mock (G-J) and sCD155-producing B16/BL6 (G and H) and CT26 (I and J) in mice treated with control mAb (mock (n = 7 (G), 12 (H), 11 (I) and 12 (J); anti-sCD155-B16/BL6 (n = 6 (G) and 12 (H)) or anti-sCD155-CT26 (n = 12 (I) and 11 (J)), or anti-PD-1 mAb (mock (n = 6 (G), 12 (H and I), and 11 (J); anti-sCD155-B16/BL6 (n = 6 (G) and 12 (H)) or anti-sCD155-CT26 (n = 4 (I) and 13 (J)). Data are pooled from two independent experiments (B-J). Statistical significance was determined by two-way ANOVA (B-F) and one-way ANOVA (G-J). Error bars indicate means ± SD. *p < 0.05; **p < 0.01; ***p < 0.001; ****p < 0.0001.

**Supplemental Figure 3.**
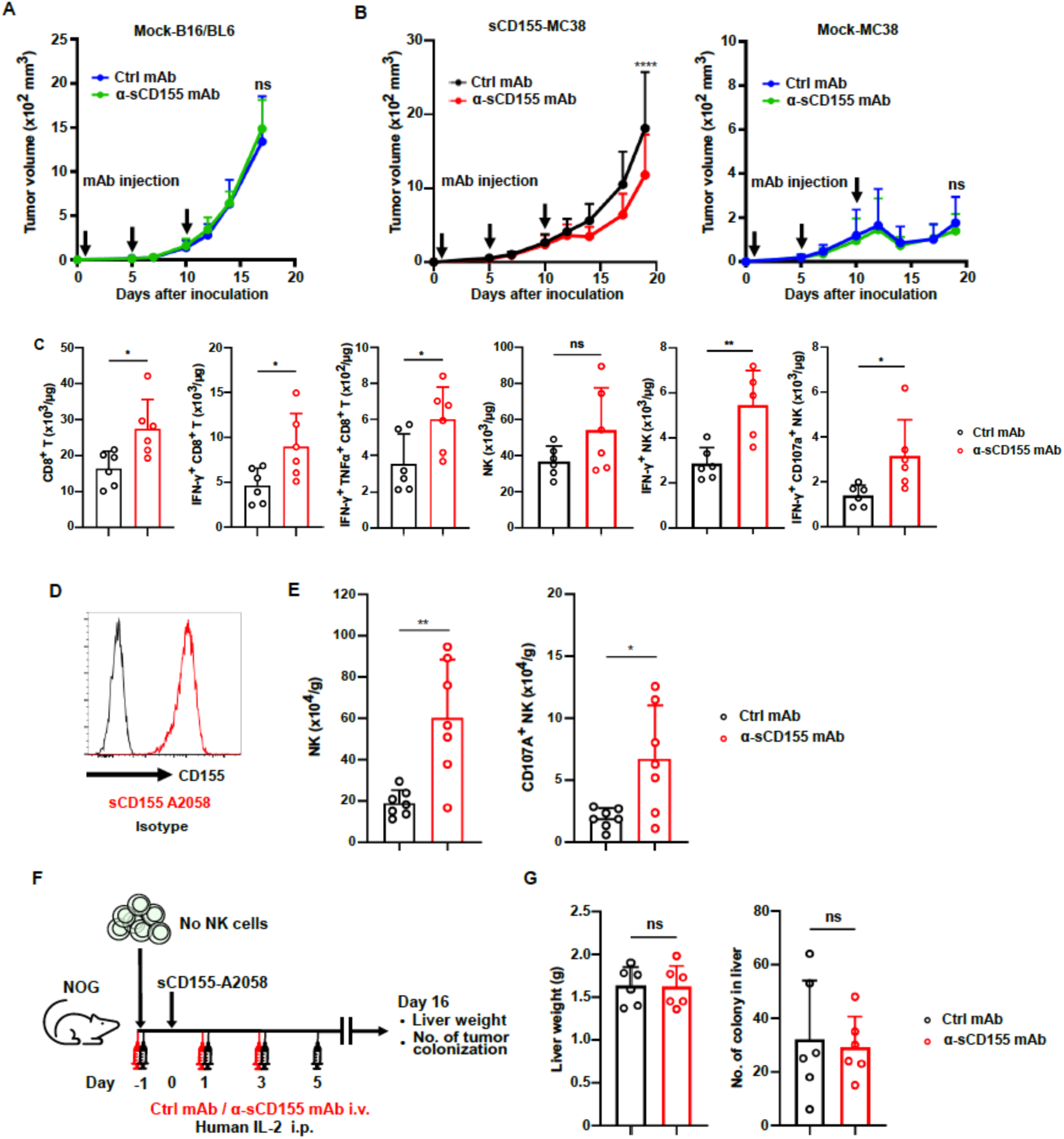
Specificity and efficacy of anti-sCD155 mAb in murine and humanized models. **(A and B)** Growth of mock-B16/BL6 in mice treated with control mAb (n = 19) or anti-sCD155 mAb (n = 19) (A), mock-MC38 and sCD155-producing MC38 (B) tumors in mice treated with control mAb (mock-MC38 (n = 8); sCD155-MC38 (n = 8)) and with anti-sCD155 mAb (mock (n = 11); sCD155-producing (10)). **(C)** Numbers of tumor-infiltrating CD8+ T cells and NK cells in sCD155-producing B16/BL6 tumors on Day 10 in mice treated with control mAb (n = 6) and anti-sCD155 mAb (n = 5). **(D)** Flow cytometry analysis of CD155 expression on sCD155-producing human melanoma A2058 cells. **(E)** Number of liver-infiltrating human NK cells in NOG mice on Day 5 post-A2058 inoculation (n = 7 per group). **(F and G)** Schematic representation of anti-sCD155 mAb treatment in the A2058 liver metastasis model (F) and quantification of liver weight and metastatic colonies (n = 6 per group) (G). Data are pooled from two independent experiments (A, B, E, G) and are representative of two independent experiments (C). Statistical analyses were performed by two-way ANOVA (A, B) or unpaired t-test (C, G). Error bars indicate means ± SD. *p < 0.05; **p < 0.01; ***p < 0.001; ****p < 0.0001.

**Supplemental Figure 4.**
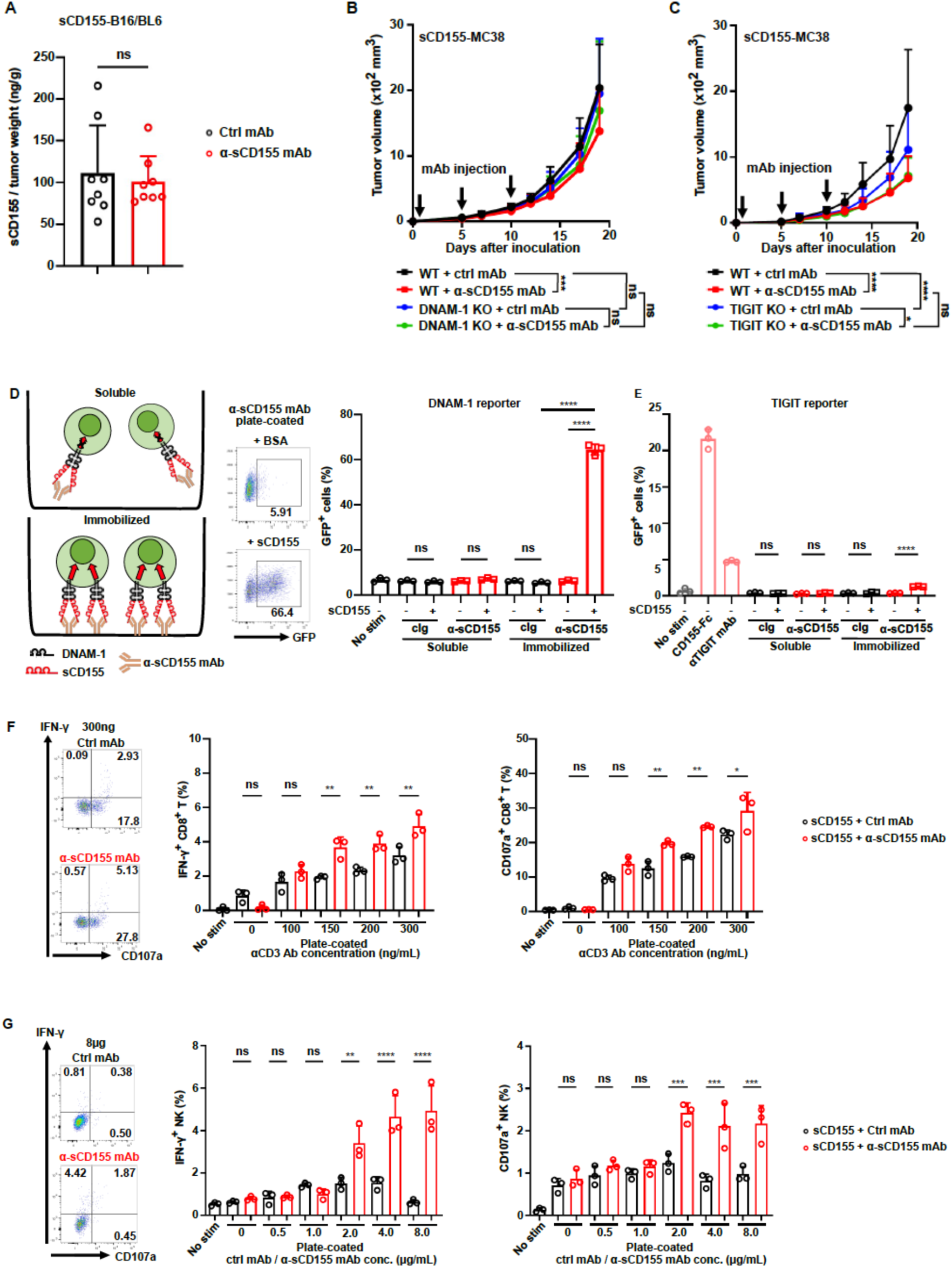
Mechanism of anti-sCD155 mAb-mediated DNAM-1 signaling and NK cell activation. **(A)** Concentration of sCD155 in tumor interstitial fluid (TIF) from B16/BL6 tumors treated with control or anti-sCD155 mAb (n = 8 per group). **(B)** Growth of sCD155-CT26 tumors in WT, DNAM-1) KO, or *Tigit* KO mice treated with control mAb (WT (n = 11 (B, C); DNAM-1 KO (11); *Tigit* KO (9)) or anti-sCD155 mAb (WT (n = 11) (B, C); DNAM-1 KO (11); *Tigit* KO (10)). **(D and E)** Schematic representation and flow cytometry analysis of reporter assays. DNAM-1 (D) or TIGIT (E) reporter cells were stimulated with soluble or immobilized control mAb or anti-sCD155 mAb in the presence or absence of sCD155) (n = 3 or 4 per group). Plate-coated soluble chimeric protein consisting of the extracellular portion of CD155 and the Fc portion of human IgG1 and anti-TIGIT mAb was used as a positive control. (**F and G)** Activation status of tumor-infiltrating IFN-γ+ and CD107a+ CD8+ T cells (F) and NK cells (G) quantified by flow cytometry (N = 3 per group). Data are pooled from two independent experiments (A-C) and are representative of two independent experiments (D-G). Statistical analyses were performed by unpaired t-test (A), two-way ANOVA (B and C), and one-way ANOVA (D-G). Error bars indicate means ± SD. *p < 0.05; **p < 0.01; ***p < 0.001; ****p < 0.0001.

**Table S1.** Baseline clinicopathological characteristics of patients according to serum sCD155 level.

| Characteristic | sCD155 high<br>(n = 14) | sCD155 low<br>(n = 35) | Total<br>(n = 49) | P value |
| --- | --- | --- | --- | --- |
| **Age** |  |  |  |  |
| Median (range), years | 68.86 (52–83) | 68.6 (50–80) | 68.67 (50–83) | 0.9117 |
| **Sex, n (%)** |  |  |  |  |
| Female | 2 (14.3) | 6 (17.1) | 8 (16.3) | >0.9999 |
| Male | 12 (85.7) | 29 (82.9) | 41 (83.7) |  |
| **Smoking status, n (%)** |  |  |  |  |
| Current smoker | 5 (35.7) | 17 (48.6) | 22 (44.9) | >0.9999 |
| Former smoker | 9 (64.3) | 16 (45.7) | 25 (51.0) |  |
| Never smoker | 0 (0.0) | 2 (5.7) | 2 (4.1) |  |
| **ECOG performance status, n (%)** |  |  |  |  |
| 0 | 4 (28.6) | 12 (34.3) | 16 (32.7) | >0.9999 |
| 1 | 9 (64.3) | 21 (60.0) | 30 (61.2) |  |
| 2 | 1 (7.1) | 2 (5.7) | 3 (6.1) |  |
| **Histology, n (%)** |  |  |  |  |
| Adenocarcinoma | 11 (78.6) | 25 (71.4) | 35 (71.4) | >0.9999 |
| Squamous cell carcinoma | 2 (14.3) | 7 (20.0) | 11 (22.4) |  |
| NSCLC, not otherwise specified | 1 (7.1) | 3 (8.6) | 4 (8.2) |  |
| **Treatment type, n (%)** |  |  |  |  |
| Pembrolizumab monotherapy | 4 (28.6) | 8 (22.9) | 12 (24.5) | 0.7214 |
| Pembrolizumab + chemotherapy | 10 (71.4) | 27 (77.1) | 37 (75.5) |  |
| Carboplatin + pemetrexed | 8 (57.1) | 20 (57.1) | 28 (57.1) | >0.9999 |
| Carboplatin + paclitaxel | 2 (14.3) | 7 (20.0) | 9 (18.4) |  |
| **Best response by RECIST v1.1, n (%)** |  |  |  |  |
| Partial response (PR) | 8 (57.1) | 18 (51.4) | 26 (53.1) | 0.7612 |
| Stable disease (SD) | 5 (35.8) | 13 (37.2) | 18 (36.7) |  |
| Progressive disease (PD) | 1 (7.1) | 4 (11.4) | 5 (10.2) |  |
| **Clinical stage, n (%)** |  |  |  |  |
| Stage IIIB | 0 (0.0) | 4 (11.4) | 4 (8.2) | 0.6561 |
| Stage IIIC | 1 (7.1) | 2 (5.7) | 3 (6.1) |  |
| Stage IVA | 6 (42.9) | 12 (34.3) | 18 (36.7) |  |
| Stage IVB | 7 (50.0) | 17 (48.6) | 24 (49.0) |  |
| **Tumor size, n (%)** |  |  |  |  |
| T4 | 9 (64.4) | 17 (48.6) | 26 (53.1) | 0.1035 |
| T3 | 3 (21.4) | 4 (11.4) | 7 (14.3) |  |
| T2 | 1 (7.1) | 9 (25.7) | 10 (20.4) |  |
| T1 | 1 (7.1) | 5 (14.3) | 6 (11.3) |  |
| **PD-L1 expression, n (%)** |  |  |  |  |
| ≥50% | 6 (42.9) | 11 (31.4) | 17 (34.7) | 0.5156 |
| <50% | 8 (57.1) | 24 (68.6) | 32 (65.3) |  |
| **Mutation status** |  |  |  |  |
| **TP53, n (%)** |  |  |  |  |
| Positive | 1 (7.1) | 4 (11.4) | 5 (10.2) | >0.9999 |
| Negative | 13 (92.9) | 31 (88.6) | 44 (89.8) |  |
| **KRAS, n (%)** |  |  |  |  |
| Positive | 2 (14.3) | 8 (22.9) | 10 (20.4) | 0.7021 |
| Negative | 12 (85.7) | 27 (77.1) | 39 (79.6) |  |
| **FGFR1 amplification, n (%)** |  |  |  |  |
| Positive | 1 (7.1) | 2 (5.7) | 3 (6.1) | >0.9999 |
| Negative | 13 (92.9) | 33 (94.3) | 46 (93.9) |  |
| **Progression-free survival** |  |  |  |  |
| Median (range), months | 15.1 (4.9–25.1) | 17.8 (1.3–41.4) | 17.0 (1.3–41.4) | 0.0263 |
| Median survival, months | 13.6 | Undefined | — |  |
| **Serum sCD155 concentration** |  |  |  |  |
| Median (range), ng/mL | 306.4 (267.6–412.8) | 189.2 (112.8–256.7) | 222.7 (112.8–412.8) | <0.0001 |

**Abbreviations:** ECOG PS, Eastern Cooperative Oncology Group performance status; NSCLC, non-small cell lung cancer; PD-L1, programmed death-ligand 1; RECIST, Response Evaluation Criteria in Solid Tumors.

